# Bacterial Communities and Soil Interactions in Sifra and Mondial Potatoes: Pathways to Sustainable Potato Production

**DOI:** 10.64898/2025.12.28.696611

**Authors:** Varaidzo W. Chakafana, Joanna F. Dames

**Author notes:** Correspondence: Joanna Dames.

## Abstract

Plant growth promoting rhizobacteria (PGPRs) play significant roles in improving plant growth. Likewise, Mycorrhizal Helper Bacteria (MHB) enhance AM fungi fitness by promoting spore germination and providing protection against pathogens. This PGPR and MHB synergistic association can be exploited to introduce inocula that utilize bacteria for sustainable potato production within potato agricultural systems. In this study, we isolated and identified PGPRs that dominate Sifra and Mondial potato soils across 3 developmental stages (sprouting, tuber bulking and at harvest). In addition, soil physicochemical properties were assessed, revealing soil pH as a key factor shaping bacterial community composition. Community structure in both soil samples was dominated by the phyla Actinobacteria, Proteobacteria, and Acidobacteria. The genera *Sphingomonas* and the families *Conexibacteraceae* and *Cthoniobacteraceae* were consistently abundant across all growth stages in both potato varieties, suggesting that these taxa represent dominant bacterial groups that are adapted to the acidic soils associated with Sifra and Mondial cultivation. Functional characterization identified nitrogen supplying bacterial species in Sifra soils while Mondial soils were enriched in biopolymer degrading bacteria reflecting a varietal influence of bacterial communities. Bacterial community was not affected by growth stage. To identify MHB, isolated bacteria from spore surfaces and within spores were cultured, compared using 16S rRNA gene sequencing, and tested for chitinase, protease, cellulase and phosphatase activity. MHB from the Mondial field were dominated by *Sphingobium* whilst those from the Sifra field were dominated by *Bacillus* species suggesting varietal influence of MHB associating with prevailing AM fungi species. All *bacillus* species facilitated phosphate solubilization, Indole Actic Acid (IAA) production and cellulase activity suggesting their roles in plant growth. Similarly, Mondial MHB had the same activity. Overall, the study highlighted varietal and environmental influence of plant PGPRs and MHB which can be exploited for sustainable potato production in South Africa.

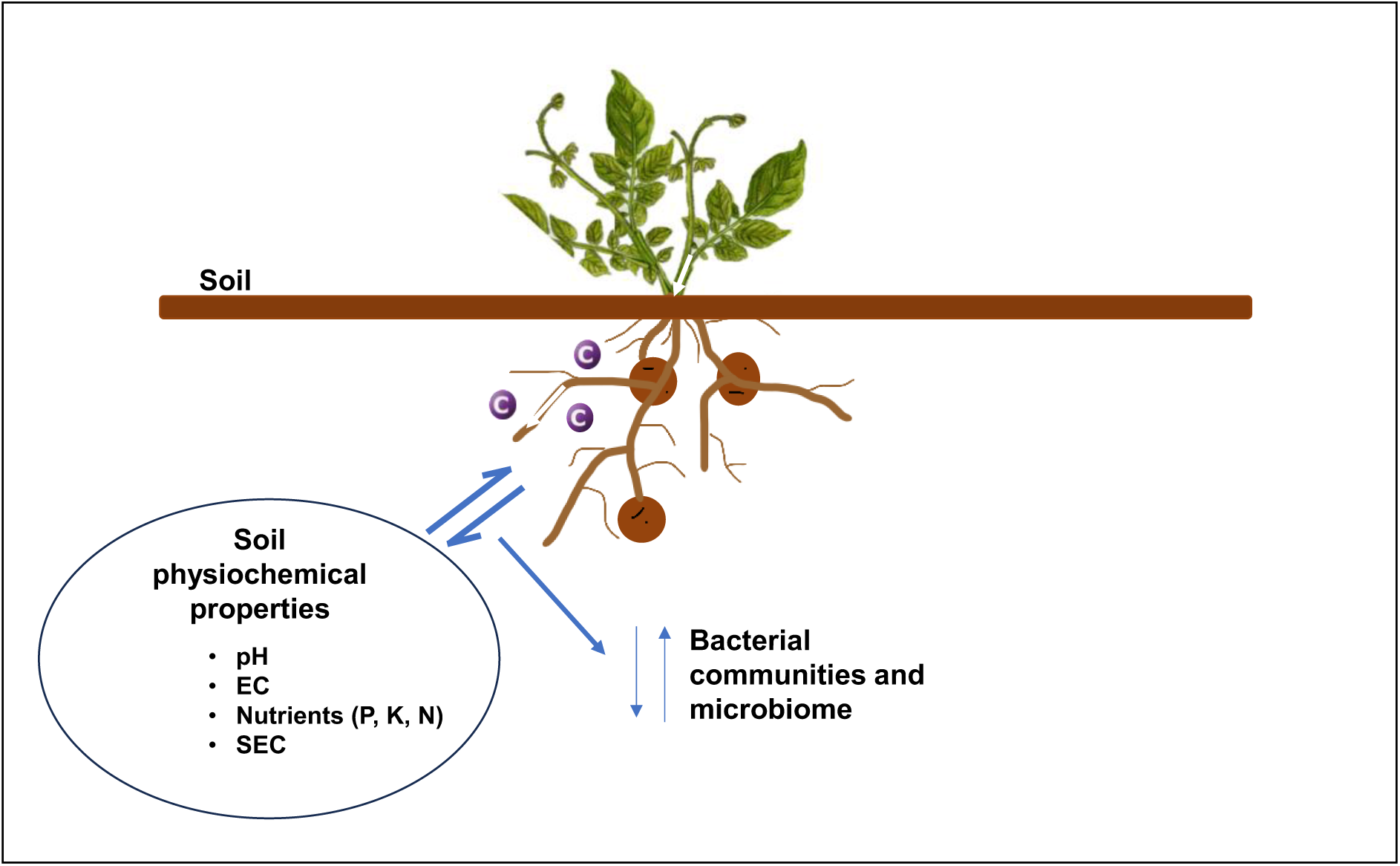

**Highlights:**

- Soil physiochemical properties are affected by potato variety
- Interaction of plant variety and soil physicochemical properties influences plant microbiomes
- Mycorrhizal helper bacteria can possess plant growth promoting properties
- Plant growth stage does not necessarily influence rhizhospheric bacterial communities

## 1. Introduction

*Solanum tuberosum* (the potato) is the most important vegetable grown in the world [1]. Approximately 375 million tons of potatoes were grown in 2022 across the world (FAOSTAT, 2024). Potatoes provide carbohydrates, vitamins and nutrients vital for human health. Most importantly is the provision of antioxidants which reduce the incidence of cancer, liver and heart disease [2]. In South Africa, potatoes are grown across 16 ecological regions [3] and are the largest fresh vegetable crop [4]. They support the livelihoods of both small- and large-scale farmers and contribute to the country’s foreign exchange earnings. They are popular for their use in French fry production. High production is met by heavy injections of synthetic fertilizers and pesticides to meet potato demands. Synthetic fertilizers such as phosphates are nonrenewable which has led to increased costs [5]. Heavy injections of macronutrients without the addition of organic amendments, secondary nutrients and micronutrients have reduced soil fertility [6]. Climate change has led to increased temperatures which have seen increased proliferation of pests and diseases. Consequently, potato prices are skyrocketing with a 100% price increase reported in 2024 [7]. Potatoes also have a short root system yet demand high water supply. Water makes up 90–95% of the potato plant tissue and 70–85% of tubers [4], making it heavily dependent on rainfall and irrigation [8]. This has led to concerns about the water balance for human consumption, industry and farming consequently leading to increased water bills [7]. This is costly for both farmers and consumers at all levels.

The advent of technology has allowed microbial characterization and investigation of their role in the ecosystem. Many ecologists argue that the soil has sustained itself and supported vegetation growth long before human intervention [9]. Agriculture can rely on soil microbes for soil improvement and plant growth and development. The root tip exudes 30% of photosynthesis that attracts 10 – 1000 times higher bacteria than bulk soil [10, 11]. These bacteria include Plant growth promoting rhizobacteria (PGPR). The use of PGPR is an effective approach to grow potatoes without the use of synthetic fertilizers and pesticides. Directly or indirectly, PGPR improve plant nutrient and water uptake [12], produce hormones which regulate plant growth and provide tolerance to biotic and abiotic stress. They do this by modifying root morphology which increases permeability to water and nutrient uptake [12]. Several reports are available of the beneficial effects of PGPRs in potato soils. Bacteria *P. agglomerans* strain P5, *Microbacterium laevaniformans* strain P7, and P. *putida* strain P13 have been involved in phosphate solubilization for potato growth [13]. *Chryseobacterium* C138, *Bradyrhizobium* and *Rhizobium* were identified as producing siderophores which are transported to plants for iron uptake [14], [15]. Bacillus species producing Indole Acetic acid (IAA) were reported to induce potato growth and development [16]. Potato PGPR strains PRP-6, PRS-17 and PRS- 24 were identified as capable of producing IAA during potato production [17]. *Achromobacter xylosoxidans* Cm4, *Pseudomonas oryzihabitans* Ep4, and *Variovorax paradoxus* C2 were reported as producing auxins and ACC deaminase which reduced amino acid exudation into the rhizosphere translating to a 40% increase of potato tuber weight [18]. PGPR antioxidants which include catalase (CAT), peroxidase (POD) and superoxide dismutase (SOD) relieve potato plants from oxidative stress [19] under drought and salinity conditions. Abscisic acid (ABA) production improves water conductivity and reduces water loss [20, 21]. *Methylobacteria* sp have been shown to increase potato health under salt stress [22]. Plant growth-promoting rhizobacteria (PGPR) can suppress potato pathogens, including *Fusarium oxysporum*, *Rhizoctonia solani*, and *Aspergillus niger*, through the production of antibiotics [23], [24] and lytic enzymes. The increase in potato productivity has generally been shown to correlate with the presence of plant growth-promoting rhizobacteria (PGPR) [25].

Bacteria can also be found physically attached to other beneficial microbes such as arbuscular mycorrhiza (AM) fungi. These mycorrhizal helper bacteria either live on the spore surface or survive within the spore cytoplasm as endosymbionts [26]. They provide fungal fitness and functionality which in turn improves plant growth and development. These MHB are also not necessarily limited to the spores as they can exist in the rhizosphere prior to any association [27]. Some MHB also have PGPR traits. Boussageon et al., 2023[29] identified a Bacillus consortium that synergistically interacted with AM fungi communities, improving P nutrition and yield. The addition of *Bacillus amyloliquefaciens* FZB42 and AM fungi + *Pseudomonas brassicacearum* 3Re2-7 significantly mitigated drought stress [30]. *Paenarthrobacter nicotinovorans* and *Bacillus circulans* from rhizospheric soils together with *Oidiodendron maius* 143 were found to promote each other’s growth while co-inoculation promoted the growth of blueberry seedlings [27]. *P. fluorescens* _F140_ and *P. fluorescens* _T17-4_ association with *Funneliformis mosseae* and *R. intraradices* increased plant biomass and chlorophyll content of potato plants [31].

Bacterial communities vary depending on host species, developmental stages, and environmental conditions [32]. Understanding and harnessing these interactions between plants and beneficial microorganisms offers significant potential for advancing sustainable agriculture, improving nutrient use efficiency, and enhancing food security [6]. In this study, we aimed to isolate and identify PGPR and MHB as well as to evaluate the biochemical properties of isolated MHB in Mondial and Sifra potato fields across the sprouting, tuberization, and harvest stages. We hypothesized that plant growth-promoting bacterial (PGPB) communities are differentially influenced by potato variety, developmental stage, and soil environment. Elucidating these interactions will enable the identification of beneficial PGPR associated with Sifra and Mondial potatoes that could be exploited to support sustainable potato production in South Africa. To our knowledge, this study is the first identification of bacteria that associate with Sifra and Mondial potato varieties.

## 2.0 Materials and Methods

### 2.1 Sample collection and DNA extraction

The study site was a farm located 48.2 km from Grahamstown in the Eastern Cape Province of South Africa (33°30’31"S, 26°52’05"E, elevation 226 m, as per Google Earth). The farm practices semi-conservation agriculture, using seaweed fertilizer, humates, kraal manure, organic manure, and crop rotations, following the sequence: potato, butternut, and peppers for two years, followed by four years of Rhodes grass. The research was conducted after a four-year period of continuous Rhodes grass cropping. Two potato varieties, Sifra and Mondial, were grown in separate fields, each subdivided into plots consisting of 14 rows, each 1 m wide and 14 m long. Rhizospheric soil was collected from Sifra and Mondial potato fields during three key developmental stages: tuber sprouting (GS1), tuber bulking (GS4), and potato harvest (PH). For each variety, three plots were selected across the three growth stages, with five sampling points per plot. The five samples from each plot were pooled, resulting in three replicates per variety for each growth stage. In total, 18 soil samples were analyzed. The soil’s physiochemical properties were assessed at the Eco Analytica Laboratory, North-West University, South Africa. The following parameters were assessed: oxidizable organic matter (C), calcium (Ca), magnesium (Mg), potassium (K), phosphorus (P), sodium (Na), cation exchange capacity (CEC), soil pH (KCl), electrical conductivity (EC), nitrogen (N), and base saturation. Bacterial DNA was extracted following the Quick-DNA™ Fecal/SoilMicrobe MiniPrep Kit (ZymoResearch Cat # D6010) according to the manufacturer’s instructions. DNA was quantified by a Nanodrop spectrophotometer (Thermo Scientific™ NanoDrop 2000). The integrity of the isolated DNA was evaluated by electrophoresis in a 1% (w/v) agarose gel at 80V for 1hr in 1x Tris/Borate/EDTA (TBE) buffer and visualized in a Chemi Doc™ XRS+ System with Image Lab™ Software.

### 2.2 Sequencing

The small ribosomal subunit (16S rRNA) was targeted to amplify the V4-V5 hypervariable region of bacterial DNA using the Miseq 515F-Y (5′-TCGTCGGCAGCGTCAGATGTGTATAAGAGACAGGTGYCAGCMGCCGCGGTAA-3′) and Miseq 926R (5′-GTCTCGTGGGCTCGGAGATGTGTATAAGAGACAGCCGYCAATTYMTTTRAGTTT-3′) primers (Parada, Needham, and Fuhrman, 2016), with Illumina sequencing adaptors. PCR reactions, conducted in an Applied Biosystems 2720 Thermal Cycler, were performed in a 25 µL final volume consisting of 12.5 µL G2 Hot Start Green Master Mix (containing dNTPs, MgCl₂, and Taq polymerase; Promega, Ref: M7422, Lot: 0000 5248 72), 0.2 µM of each primer, and 1 µL of DNA. The PCR cycling conditions included an initial denaturation at 95^°^C for 5 minutes, followed by 30 cycles of denaturation at 95^°^C for 45 seconds, annealing for 45s at 50^°^C, extension for 1 min at 72^°^C with a final extension of 5 mins at 72^°^C and an infinity hold of 4^°^C. Amplification of the ∼500bp band was verified by running a 1% agarose gel. Successfully amplified PCR products were cleaned using the Promega Wizard R SV Gel and PCR Clean-Up System (Promega Ref: A9281, LOT: 0000 4680 22). The DNA was sent for Illumina Miseq Sequencing at the Next Generation sequencing laboratory at the South Africa Institute for Aquatic Biodiversity (SAIAB) at Rhodes University. In summary, Nextera XT indices (Illumina) were added for multiplexing purposes via a secondary PCR step as described by the manufacturer and purified using AMPure XP beads (Beckman Coulter). Prepared libraries were quantified with Qubit dsDNA HS assay (Invitrogen) on a Qubit™ 4 Fluorometer (Qubit, ThermoFisher Scientific, South Africa) and the quality of the pooled libraries verified using the Bionalyser HS dsDNA chip (Agilent) on the Bioanalyser 2100 platform (Agilent, Merck, South Africa). Successful addition of multiplex adaptors was confirmed by qPCR using NEBNext Library Quant Kit for Illumina (NEB) on a QuantStudio 3 (Applied Biosystems, ThermoFisher Scientific, South Africa). The libraries were sequenced on a MiSeq platform (Illumina, California, USA) with Nextera v3 600 cycle (Illumina) kit with a 10% PhiX spike.

### 2.3 Sequence processing and Statistical Analyses

Raw forward and reverse reads from each sample were paired to create single files for analysis in Mothur. Ambiguous reads were removed by specifying maximum (∼550 bp) and minimum (100 bp) expected lengths. The VSEARCH algorithm was used to eliminate chimeras. Sequences were classified using the Bayesian classifier against the SILVA reference database (Release 138.1), and undesirable lineages, including Chloroplast, Mitochondria, Archaea, Eukaryota, and unknown lineages, were excluded. The remaining sequences were realigned to the SILVA reference database (silva.nr_v138_1.align) and screened to ensure consistent coverage across the V4-V5 region. To facilitate data comparison, sequences were subsampled to 32,412 sequences per sample. Operational Taxonomic Units (OTUs) were generated using a pairwise distance cutoff of 0.03 and clustered with the furthest neighbor algorithm. Mothur outputs (shared file and cons.taxonomy files) together with a generated metadata file were inputted into R (version 4.1.1 and R studio version 2022.07.1+554), MicrobiomeAnalyst and Primer6 PERMANOVA for statistical analysis with some data graphically represented using GraphPad Prism (Version 8.0.2 (263). MicrobiomeAnalyst generated rarefraction curves, calculated relative abundance and beta and alpha diversity profiling at 95% confidence intervals. It utilized the linear discriminant analysis (LDA) method that captured and discriminated biological data through differences in taxonomy, genes and functions. The effect size (LEfSe) determined the magnitude of these differences. Linear Discriminant Analysis (LDA) Effect Size (LEfSe) at feature level on a 95% confidence interval with variety as experimental factor analyzed the top 20 different features (taxonomy, genes or functions which capture differences) at LDA 1. OTUs with significant differences between Sifra and Mondial were identified. Therefore, LEfSe was used to test violation of the null hypothesis that there was no difference between biological data. Due to some limitations in MicrobiomeAnalyst, R calculated statistical significance of alpha diversities within varieties at 95% significance using data collected from MicrobiomeAnalyst. Primer6 PERMANOVA was used to conduct Principal Coordinate Analysis (PCA), dominance plots, RELATE and non-metric multidimensional scaling (MDS). Biological data was fourth root transformed [log(y+1)] as most counts were 0 whilst environmental data was normalized. Principle Component Analysis (PCA) was used to identify relationships and groupings of samples based on their environmental data. Both transformed biological and environmental data were used to generate resemblance matrixes using the Bray-Curtis and Elucidian coefficients respectively. Generated resemblance matrices were used to RELATE the biological data to the environment at a 95% significance level. MDS and PICO plots for both biological and environmental factors were generated. For taxonomic assignment, a representative sequence from each OTU generated in the Mothur FASTA file was selected for BLASTn search. Ten hits from each representative sequence were saved. In this analysis, only those with a query cover above 90% and percentage identity of above 95% were considered for analysis unless specified. Assignment to different taxonomic levels depended on query cover and percentage. Percentage identity of 61.39 classified Kingdom, Phylum - 69.30, Class - 75.39, Order - 83.40, Family - 90.95, Genus - 93.07, Species - 98.62 and Strain - 99.59. Mega II was used to generate phylogentic trees of varying bacterial species using the neighbor-joining algorithm.

### 2.4 Mycorrhizal Helper Bacteria (MHB) identification

#### MHB isolation

Arbuscular mycorrhizal (AM) fungal spores were extracted using the sucrose density gradient centrifugation method according to [33], [34]. Briefly, for each sample, soil from each trap culture pot was mixed with approximately 300 mL of water in a beaker and vigorously stirred using a sterile spatula. The mixture was allowed to settle for 1 minute, and the supernatant was decanted through a series of sieves (425 μm, 250 μm, 125 μm, and 45 μm). This process was repeated four times. Debris collected in 250 μm, 125 μm, and 45 μm sieves was transferred into separate 50 mL falcon tubes, resuspended in water, and centrifuged at 1900xg for 5 minutes. The supernatant was carefully discarded, and the pellet was resuspended in 60% sucrose solution, then centrifuged again at 1900xg for 5 minutes. A vacuum pump (Model 801; Cat 167801-22) was set up, and 9 cm Whatman No. 1 filter papers were gridded and labeled to correspond to the falcon tubes. The supernatant from each tube was decanted onto the filter paper, and the solution was filtered, trapping the spores. The filter paper, still under suction, was rinsed with water to evenly distribute the spores. The filter paper was then transferred to a clean petri dish, covered, and left to dry. Isolated spores were transferred into 60 μL of 0.2% saline solution. Due to the low spore count, the entire volume was spread onto Tryptone Soy Agar (TSA, Art. No. HG00°C17.500) plates. The plates were incubated overnight at 28^°^C. Single colonies were then streaked discontinuously to obtain pure bacterial cultures. These cultures were subsequently grown in Merck BioLAB Nutrient Broth (NB) media (Biolab, Art. No. HG00°C24.500) and incubated at 37^°^C for 5 days at 200 rpm to support the growth of slow-growing bacteria.

Bacteria was pelleted by centrifugation of broth cultures at 15000xg for 5 mins and resuspended in 200ul bashing bead buffer. DNA was isolated using the Quick-DNA™ Fungal/Bacterial Miniprep Kit (ZymoResearch, Cat # D6005) according to the manufacturer’s instructions. DNA was measured by a Nanodrop spectrophotometer (Thermo Scientific™ NanoDrop 2000). Integrity of the isolated DNA was evaluated by electrophoresis in a 1% (w/v) agarose (Anatech K&K Scientific, 923866) gel at 80V for 1hr in 1x Tris/Borate/EDTA (TBE) buffer and visualized in a Chemi Doc™ XRS+ System with Image Lab™ Software. The PCR reactions (25 μl final volume) were carried in an Applied Biosystems 2720 Thermal Cyclerand comprised of 12.5 μl GoTaq Hot Start Green Master Mix (Promega Ref:M7422, Lot: 0000 524872)), 0.2μM of each primer and 5μl DNA template. The PCR amplified the 16S RNA region using the general fD1 (5’-GAGTTTGATCCTGGCTCA-3’) and rP2 (5’-ACGGCTAACTTGTTACGACT-3’) bacterial primers (Weisburg et al., 1991). The PCR cycling conditions included an initial denaturation of 5 mins at 95°C followed by 30 cycles of denaturation for 45 s at 95^°^C, annealing for 45 s at 50^°^C and extension for 1 min at 72^°^C with a final extension of 5 mins at 72^°^C and an infinity hold of 4^°^C. Amplification of the ∼1500bp band was verified by running a 1% agarose gel. Successfully amplified PCR products were cleaned using the Wizard R SV Gel and PCR Clean-Up System (Promega Ref: A9281). The samples were sequenced at Inqaba Biotechnical Industries (Pty) Ltd using the sanger sequencing technique.

### 2.5 MHB biochemical properties

#### 2.5.1 Phosphate solubilization

The ability of MHB to solubilize phosphate was assessed using a modified version of the National Botanical Research Institute’s Phosphate (NBRIP) growth medium (Table S1), with bromophenol blue as an indicator. Agar discs were removed from the NBRIP plates, and 100 µL of MHB culture was placed into the wells then incubated at 28°C. Phosphatase production, indicating phosphate solubilization, was evidenced by a blue coloration around the edges of the wells. Further solubilization was indicated by the formation of a clear zone around the bacterial culture as the pH decreased.

#### 2.5.2 Chitinase production

Bacterial isolates were grown in LB broth overnight with constant shaking at 36°C then streaked onto chitin agar (Table S2) and incubated at 37°C for 3 days. Bacterial growth on the chitin agar indicated chitinase production, as chitin served as an alternative carbon source.

#### 2.5.3 Cellulase production

Cellulase production was evaluated by removing discs from carboxymethylcellulose (CMC) agar (Table S3) and filling the wells with 100 µL of the bacterial isolate growing in LB broth. The plates were incubated at 30°C for 5 days. Following incubation, 5 mL of iodine solution was added to the plates; iodine binds to undegraded cellulose, while a yellow coloration in the zones of clearance indicated cellulase degradation

#### 2.5.4 Protease production

To test the ability of mycorrhiza helper bacteria (MHB) to produce proteases for spore degradation, skimmed milk agar was used. Disc wells were created in the agar, and 100 μL of bacterial solution was added to each well. The plates were then incubated at 28°C for 5 days. The appearance of halos around the wells indicated protease production, as the proteases degraded the milk proteins in the agar.

#### 2.5.5 Production of Indole Acetic Acid

The IAA test was performed by incubating bacterial isolates in 1ml of Nutrient broth with shaking at 200rpm overnight. A 100ul of incubated bacterial solution was placed in 1ml of DEV Trptophan (Merck Cat# 1.10694.0500) broth and incubated for 5 days with 200rpm continuous shaking at 37^°^C. The cells were centrifuged at 15000rpm for 20mins and 300ul supernatant collected. Six hundred microlitres of Kovacs Reagent (MERCK Cat# SAAR3745000JT) was added to the supernatant. IAA production was confirmed by colour change to pink and red was indicative of IAA production.

## 3.0 Results

### 3.1 Isolation and identification of bacteria in potato rhizosphere

This study employed Illumina sequencing for bacterial characterization. However, successful sequencing requires amplification of the correct genomic region. The ∼500 bp V4-V5 16S ribosomal RNA region was successfully amplified by PCR as represented by Figs. 1A and B. DNA sequencing of the amplified region generated 454 927 unique sequences which were classified and clustered into Operational Taxonomic Units at a 97% cutoff using the Furthest algorithm method. Bacterial relative abundance in both Mondial and Sifra soils identified a dominance of Actinobacteria >Proteobacteria > Acidobacteria regardless of variety. Sifra had 32.8% Actinobcteria, 25.8% Proteobacteria and 10.8% of Acidobacteria in relative abundance while Mondial had Actinobacteria (29.2%), Proteobacteria (29.1%) and Acidobacteria (11.4%) (Fig. 1C). This suggests these as important in Mondial and Sifra’s growth and development. Other phyla which were not as dominant as the first 3 included Firmicutes > Plantomucetota > Bacteroidota > Verrucomicrobiota > Chloroflexi > Gemmatimanadota > Myxococcota in that order (Fig. 1C). After removing unclassified genera and considering the top 7 genera, similar observations were made when genera relative abundances were compared. Mondial and Sifra had a domination of *Sphingomonas*, *f_Conexibacteraceae*, *f_Chthoniobacteraceae*, *Norcardioides*, *Solirubrobacter*, *Pseudolabrys* and *KD4_96_ge* in their rhizospheres (Fig. 1D). Across Sifra’s development, *f_Chthoniobacteraceae* (5.4%) dominated at GS1, with *Sphingomonas* (5.4%) dominating at tuber bulking then a dominance of *f_Chthoniobacteraceae* (4.0%) at harvest. Similarly, Mondial’s GS1 was dominated with f_Chthoniobacteraceae while GS4 was dominated by *Sphingomonas* though at a lower abundance than Sifra (4.1% and 3.8 respectively). Contrary to Sifra’s *f_Chthoniobacteraceae* domination at harvest, Mondial’s harvest was dominated by *Sphingomonas* at (3.6%).

**Fig 1.**
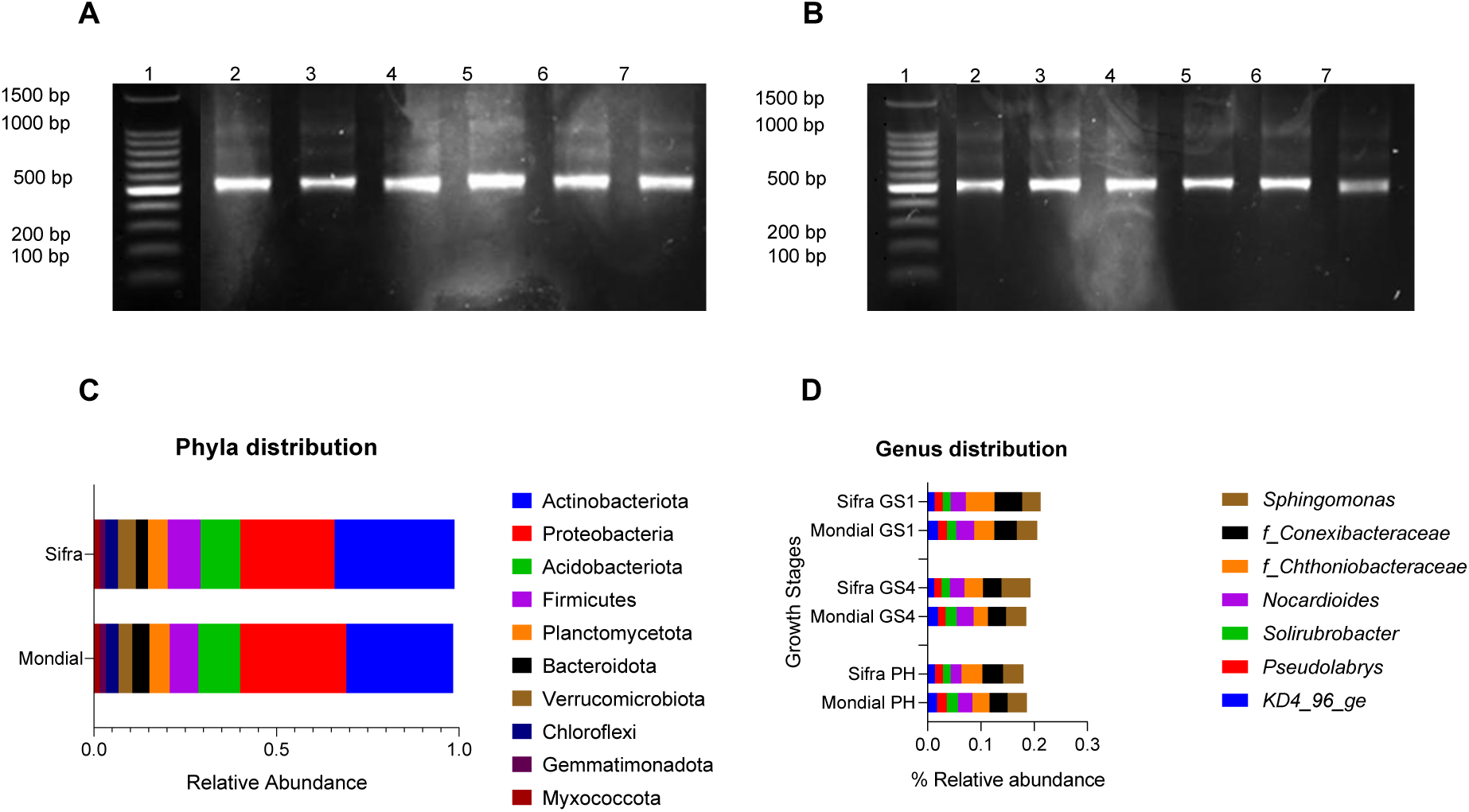
Bacterial molecular characterization. A - Representative image of bacterial DNA in Mondial soil samples showing the expected ∼500bp band. B – Representative image of bacteria in Sifra soil samples with expected ∼500bp band. C – Bacterial phyla relative abundances in Mondial and Sifra potato soils. D – bacterial genus distributions across Mondial and Sifra’s growth stages. GS1 – Growth Stage 1 characterized by sprouting, GS4 – Growth Stage 4 characterized by tuber bulking and PH – potato harvest. Similar genus dominance across development is observed in both varieties.

Despite both varieties having similar genera, a beta analysis of the samples confirmed the degree of dissimilarity. Analysis of similarities (ANOSIM) utilizing the PCAo ordination method confirmed the significant difference at feature level with R: 0.66358 and p < 0.001 (Fig. 2A) which was limited to varietal level and not the growth stage. PCAo accounted for 56% of the difference found between varieties (PCo Axis 1 accounted for 41.9% of total variation and Axis 2 accounted for 14.8%). This was confirmed by the Linear Discriminant Analysis (LDA). Linear Discriminant Analysis identifies organismal variables and uses these to discriminate between species. LDA Effect Size (LEfSe) at feature level on a 95% confidence interval with variety as experimental factor analyzed the top 20 different features (taxonomy, genes or functions which capture differences) at LDA 4 (Fig. 2B). Further analysis of growth stages within each variety did not yield any significant differences (Mondial: R = 0.004; p = 0.464 and Sifra: R= -0.235 and p = 0.836) (Fig. 2B). The analysis of variance between the samples were further analyzed utilizing cluster analysis of biological data (Fig. 2C). The samples grouped according to variety indicating a varietal influence of bacterial species except for samples S2, S5 and S8 which were closely related to the Mondial samples.

**Fig. 2.**
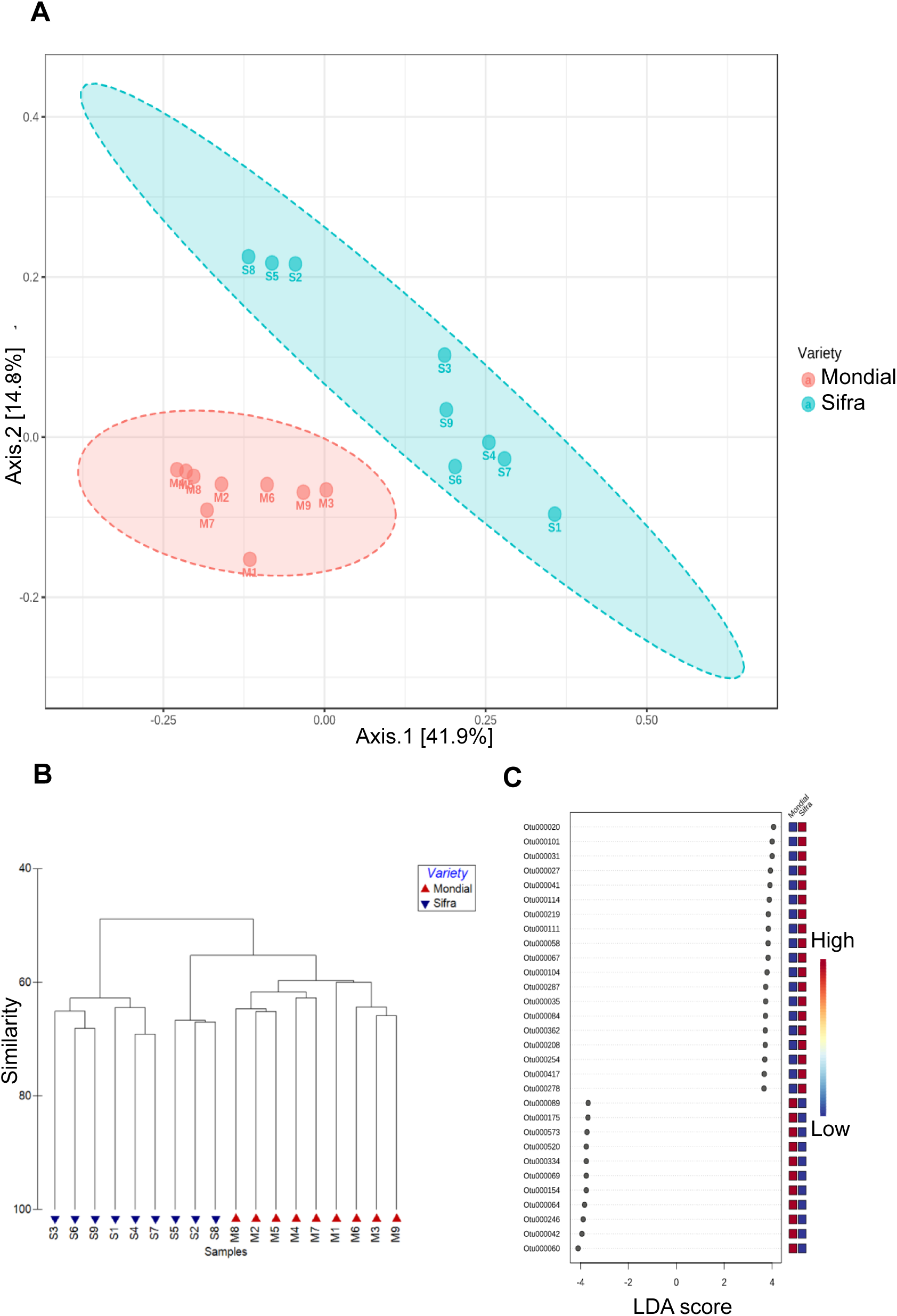
Bacterial species patterns in soil samples. A - Principal coordinate analysis (PCoA) ordination of a Bray-Curtis dissimilarity matrix between the 2 varieties showing distinct bacterial grouping by variety. B – Linear Discriminant Analysis (LDA) Effect Size (LEfSe) at feature level. These features include taxonomy, genes or functions, identifying differences between Sifra and Mondial bacterial diversities. C – Hierarchical clustering of Mondial and Sifra samples showing phylogenic distinctions between the varieties.

### 3.2 Diversity of bacterial compositions in Sifra and Mondial soils

Alpha diversity of bacterial communities was assessed using three ecological indices: Chao1 (species richness), Shannon (diversity), and Simpson (dominance). Mondial had significantly higher species richness and diversity (Chao1 index - 2840; Shannon index - 7.44 respectively) compared to Sifra (Chao1 index - 2672, Shannon index - 7,29) (Fig. 3A) with Chao1 p-value: 0.0083132; [T-test] statistic: 3.1526 and Shannon index p-value: 0.011254; [T-test] statistic: 2.8682) between the 2 varieties. In contrast, no significant difference at feature level was observed when dominance (Simpson index) was assessed between the 2 varieties (p= 0.2629) (Fig. 3C). Despite this, site analysis identified M3, M6 and M9 as having the highest number of bacterial species (Supplementary data, Fig.1 S1.). The high species number automatically made Mondial more diverse than Sifra as indicated by the Shannon index (7.44 versus 7.29). The same soil that harbored highly diverse AM fungi also harbored equally highly diverse bacteria suggesting a correlation between bacterial species and AM fungal species. Analysis within varieties across the 3 developmental stages showed no significant differences. Simpson, Shannon and Chao1 indices for Mondial had p= 0.8752; 0.1849 and 0.7326 respectively. The same test for Sifra developmental stages were p = 0.1931; 0.1849 and 0.7326 respectively. These nonsignificant differences were identified when either parametic 2-way ANOVA or non-parametric Kruskal Wallis was carried out.

**Fig. 3.**
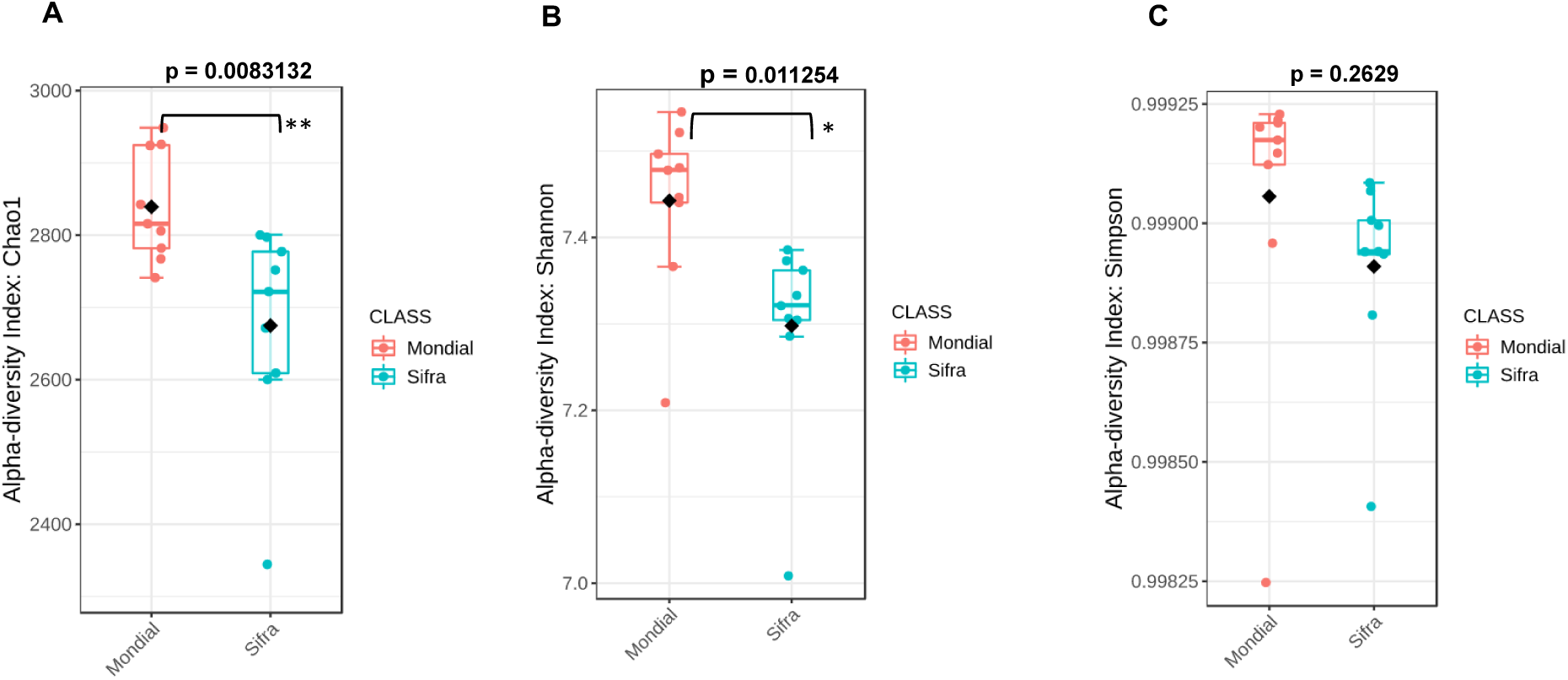
Alpha diversity of the soil bacterial community according to Chao1, Shannon and Simpson at feature level respectively. T-test was used to compare diversity between varieties. P – probability value. ***p < 0.001; **p < 0.01; *p < 0.05.

### 3.3 Environmental influence on biological data and bacterial functional prediction

We then explored the relationship between the soil physiochemical properties and biological data by understanding the variables in the soil samples. Principle Component Analysis (PCA) showed PC1 as negatively correlated with Available Na, Available K, CEC, EC and % Carbon (Fig. 4A). This means a decrease in these variables would increase the PC1 value. Sifra samples were characterized by these properties suggesting a higher presence of Available Na, Available K, CEC, EC and % Carbon (Fig. 4A). PC2 was negatively correlated with base saturation % and pH(KCl). This means a decrease in these values increased the PC2 value. Mondial soil samples were characterized by increased pH and base saturation% but negatively correlated with available Na, available K, CEC, EC and % Carbon. This suggests Mondial soil samples as having a higher pH compared to Sifra soil samples. It also suggests that the variation patterns of the soil bacterial community within or between Mondial and Sifra were determined by these variables (Rho 0.726, p = 001) concurring with an environmental PERMANOVA that identified sample separation by variety (pseudo-F = 6.6462, 0.02) and not by growth stage (pseudo-F = 1.1275, p = 0.349). An overlay of the environmental and biological resemblance matrix through RELATE analysis also showed an environmental effect on bacterial taxonomy (Rho 0.474, p = 0.001). A BEST analysis revealed pH and Available K as physiochemical properties best explaining these patterns. The magnitude of influence of these factors was further analyzed using Distance based linear models (DistLM) which revealed pH as the most influential variable (p = 0.001 (Fig. 4B). The addition of other variables did not change the bacterial community dynamics. Functional analysis of bacterial species OTUs under this environment show a prevalence of nitrogen reducing (*g_Gaiella* and *f_Conexibacteraceae*), carbon utilizing (*g_Niallia* and *f_Conexibacteraceae*), and cellulose utilizing (*o_Rhodospirillales*) bacteria in Sifra soils compared to Mondial soils (Table 1). The repetitive presence of nitrogen reducing bacteria suggests a deficiency in readily absorbable N in Sifra soils for uptake. On the contrary, Mondial soil harbored diverse bacteria responsible for starch degradation (*Nicardiodes*), catalayse production (*Paraburkholderia susongesis* strain L226) and utilization cellobiose (*Rhizobium altipalni* strain BR 10423) and lactose (*f_Acidobateriaceae*) (Table 1). The presence of numerous bacterial species confirms the bacterial alpha diversity identified in (Fig. 3A).

**Fig. 4.**
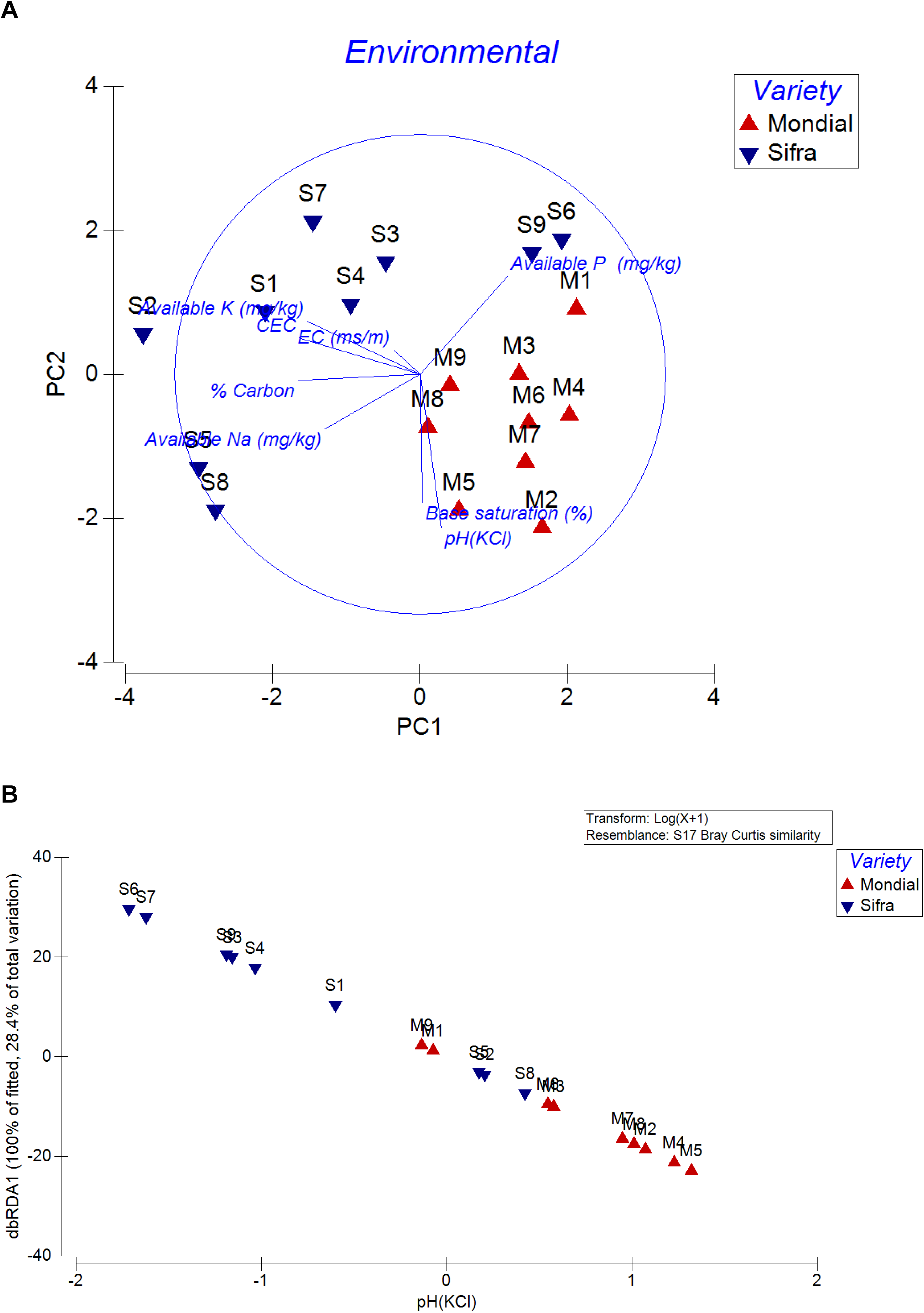
Soil physiochemical analysis. A – PCA plot of potato soil samples. B – DistLM graph showing pH as the main variable contributing to bacterial composition.

**Table 1.** Functional prediction of bacterial species identified in Mondial and Sifra soils.

| OTU | NCBI Blast result | Function |
| --- | --- | --- |
| SIFRA |  |  |
| 20, 31 | *g_Niallia* | metabolism (breakdown) of lactose, sucrose, maltose, D-mannitol, D-xylose; production of alkaline phosphatase, catalase, alpha and beta glucosidase |
| 35, 58, 114, 219, 362, 417 | *g_Gaiella* | nitrate reduction, gelatine degradation, alkaline and acid phosphatase; d-glucose, d-fructose, d-mannose utilization |
| 84, 101 | f_Conexibacteraceae | utilizes nitrate, trehalose, rhamnose, glucose; acid and alkaline phosphatase, catalase, gelatinase production |
| 27, 41 | f_Chthoniobacteraceae | breakdown of organic carbon in soil (plant biomass) |
| 111, 254 | *g_Conexibacter* | utilizes nitrate, trehalose, rhamnose, glucose; acid and alkaline phosphatase, catalase, gelatinase |
| 67 | *g_Rhodoplanes* | involved in biotin biosynthesis, gallate degradation, cyanate degradation |
| 104 | *g_Methylocapsa* | halophily, utilizes acetate, ammonium, nitrate, methane, formate, |
| 287 | *g_Chthoniobacter* | breakdown of organic carbon in soil (plant biomass) |
| 208 | *g_Methylosinus* | methane oxidation Therefore methanol dehydrogenase, methane monooxygenases, alpha-ketoglutarate dehydrogenase, nitrogen fixation, ammonia transport and assimilation, formation of nitrous oxide |
| 278 | *o_Rhodospirillales* | utilization of cellulose, carboxymethylcellulose, arabinose; produces acetoin polyhydroxybutyrate; catalase and cytochrome oxidase |
| MONDIAL |  |  |
| 89 | *Nocardioides iriomotensis* strain IR27-S3 | starch degradation, octane oxidation, glycogen metabolism, adipate degradation, glycogen metabolism |
| 175 | *g_Gemmatimonadaceae* | Adipate degradation, biotin biosynthesis, sulfopterin metabolism |
| 573 | *f_Acidobacteriaceae* | utilizes lactose, lactulose, raffinose, hydrolysis lichenan and xylan, resistant to ampicillin, kanamycin and tetracycline; alkaline phosphatase, N-Acetyl-b-glucosaminidase |
| 520 | *Hyphomicrobium* |  |
| 334 | *Paraburkholderia susongensis* strain L226 | Catalase production |
| 69 | *o_Desulfuromonadales* | synthesis of biotin, adipate degradation, ethanol fermentation |
| 154 | *g_Vicinamibacter* | 2-oxoglutarate, acetoin, alanine, casamino acids, cellobiose; sensitive only to streptomycin and not gentamycin; produces indole, acid and alkaline phosphatase, alpha-chymotrypsin |
| 64 | *g_Rhodoplanes* | utilizes acetate, malate, glutamate; pathways - starch degradation, biotin biosynthesis, |
| 246 | *Luteimicrobium xylanilyticum* strain W-15 | found in beetle (*Masticus raddei*) gut |
| 42 | *g_Gaiella* | nitrate reduction, gelatine degradation, alkaline and acid phosphatase; d-glucose, d-fructose, d-mannose assimilated |
| 60 | *Rhizobium altipalni* strain BR 10423 | halophily; utilizes cellobiose, D-fructose,galactose,mannitol, (-)-quinic acid; catalase, beta - galactosidase, -glucosidase; tryptophan deaminase |

### 3.4 Mycorrhizal Helper bacteria and their biochemical properties

DNA sequencing and analysis using the BLAST tool revealed that all the 5 mycorrhiza helper bacteria isolates shared high sequence identity (98.96–100%) with corresponding 16S rRNA gene sequences deposited in GenBank. Furthermore, phylogenetic reconstruction grouped these isolates into four genera forming two distinct clades (Fig. 5A). Bacterial diversity on AM fungi spores from Sifra and Mondial potato spores identified *Bacillus* and *Calidifontibacillus* strains as associated with spores in Sifra soils while Mondial soils allowed the proliferation of *Agrobacterium* and *Sphingobium* species on AM fungi spores (Fig. 5A). This suggests Bacillaceae as the dominant cultivable species from AM fungi in Sifa soils regardless of AM fungi species. Biochemical properties of these bacteria were evaluated *in vitro* for properties that are known to be essential for arbuscular mycorrhiza viability and functionality as well as for plant growth. The assays assessed phosphate solubilization (phosphatase activity), indole-3-acetic acid (IAA) production, cellulase activity, protease activity, and chitinase activity (Fig. 5B). Distinct functional profiles were observed among the isolates. *Calidifontibacillus erzurumensis* strain P2 and *Bacillus pacificus* strain MCCC 1A06182 exhibited phosphatase, IAA, and cellulase activities but lacked protease activity, while differing in chitinase production. In contrast, *Bacillus rhizoplane* strain JJ-63 failed to produce IAA but exhibited the other tested activities. Both *Agrobacterium* and *Sphingobium* isolates demonstrated phosphatase, IAA, cellulase, and chitinase activities but did not exhibit protease activity. These differences in bacterial species suggest a plant species influence of bacteria proliferating on AM fungi spore.

**Fig. 5.**
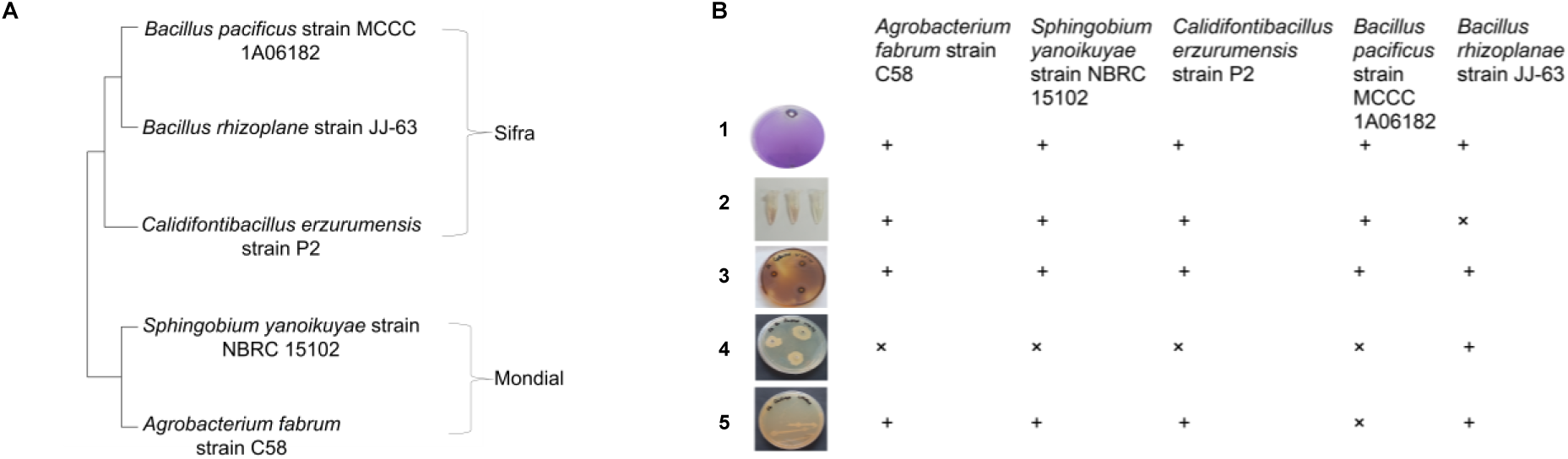
Mycorrhizal Helper Bacteria on Sifra and Mondial varietal spores. A - Neighbor-joining phylogenetic tree showing phylogenetic relationship between mycorrhizal helper bacteria isolates based on the 16S rRNA gene sequences forming 2 clades. B – Biochemical activity of identified MHB. **+** shows presence of activity and **x** shows absence of activity. B1 – phosphatase activity, B2 - IAA production, B3 - cellulose digestion B4 – protease, B5 - chitinase degradation

## 4.0 Discussion

Synthetic fertilizers and pesticides are costly, non-renewable substances that contribute to soil pollution. Plant growth promoting bacteria provide an economically and environmentally friendly solution towards sustainable plant growth with their inoculation and maintenance becoming an integral part of agriculture [8]. Isolation and identification of these prevailing PGPRs across plant development allows for their appropriate introduction/maintenance in soil towards sustainable plant development. This may also increase their interactions with other beneficial microbes. The aim of this study was to investigate potato rhizobacterial communities in South Africa towards sustainable potato production within South Africa’s potato farms. For this purpose, the most dominantly produced potato varieties, Sifra and Mondial, were chosen.

Actinobacteria > Proteobacteria > Acidobacteria > Firmicutes were found to be the most dominant phyla in Mondial and Sifra soils (Fig. 1C). Differences were observed only when abundances were compared between the 2 fields with Mondial having less abundance of Actinobacteria compared to Sifra while the opposite was true for Proteobacteria. The abundance of Acidobacteria was not very different between the 2 varieties. This is consistent with Aloo, Mbega and Makumba, 2020[35] and Purwantisari et al., 2023[36] who identified a dominance of Proteobacteria in potato rhizospheres. Similar studies identified Proteobacteria, Actinobacteria in addition to Chlorofexi as dominating potato rhizospheres [36]–[42]. It is also well known that plant exudates vary according to cultivar but the presence of similarly dominating phyla suggests them as core in potato production. Core microbiomes have been described as genomic attributes of a specific host [43]. Our results are in accordance with other studies documenting a presence of core microbiome that supports potato development [32, 33, 39, 40]. Not only are these phyla dominant in the rhizosphere but also phytosphere of potato plants [46]. However, a specific dominance of Actinobacteria, Proteobacteria and Acidobacteria in Mondial and Sifra soils without a great dominance of Chlorofexi and Bacteroidetes as in other studies suggests these as core in Mondial and Sifra’s growth and development [33, 42]. It should also be considered that many genera fall under one phylum. For instance, Actinobacteria has over 250 genera with 3000 species [48]. When bacterial genera were compared, both fields had a domination of *Sphingomonas*, *f_Conexibacteraceae*, *f_Chthoniobacteraceae*, *Nocardioides*, *Solirubrobacter*, *Pseudolabrys* and *KD4_96_ge*.

*Sphingomonas* and *Bacillus* have been described as dominant plant growth promoting rhizobacteria (PGPRs) in potato soils which supports their presence in this study though with *bacillus* at lower abundances. They can be found even in healthy soils [25, 44] and in all potato tissues at genus level [46] with farming practice increasing their abundance [50]. *Sphingomonas* were identified as increasing in abundance particularly in sludge fertilization [51]. Samples utilized in this study were from fields utilizing a semi conservative farming practice with an addition of humates. It is possible that this may have contributed to an increase in the abundance of this genus. Unlike dynamic microbiomes, core microbiomes represent the first line of defense in the rhizosphere [43]. *Bacillus* species have been implicated in disease resistance (such as *Rolstania solanacearum, Fusarium sp.* and *P. infestans* infection), and the provision of phytohormomes and metabolites for plant growth [52], [53] [36]. *Sphingomonas* provide plant stress tolerance and growth [54]. The Shingomonas was also reported to target Candida. albicans [55]. Other dominating genera such as *f_Conexibacteraceae* and *Solirubrobacter* are known for nitrate reduction while *f_Chthoniobacteraceae* are for carbon breakdown [56]. Nocardioides appear in low nutrient concentrations, degrade pollutants [57], and reduce *Staphylococcus aureus* proliferation [55] and are also involved in nitrogen reduction. Pseudolabrys is involved in nitrogen fixation [50] while the genus KD4_96_ge contains species found in contaminated soils [58]. The presence of these genera in this study suggests a nutrient provision, tolerance to biotic and abiotic stress and plant growth to our varieties. This is important as Sifra and Mondial are susceptible to disease. Sifra is highly susceptible to late blight and common scab [59] while Mondial is susceptible to Potato Virus Y (PVY) and powdery scab [59]. The genera identified in this study greatly differ with other studies identifying *Actinomycetes* and *Pseudomonas* as dominating potato rhizospheres compared to their roots [44] with rhizospheric *Azobacter* and *Bacillus* populations increasing with potato growth [56, 57]. The differences in genera observed between the current study and others suggest that as the taxonomy levels reduce, there begins a differential association of bacterial species in accordance to the interaction between the local environment and vegetation. This study represents the first to identify Conexibacteraceae and *f_Chthoniobacteraceae* as being associated with potatoes.

Our study investigated the effect of plant growth stages on potato rhizospheric bacteria by assessing bacterial community abundance and richness. Bacterial diversity has been reported to change from the seedling stage to harvest in potato soils [42]. Our study showed differences in abundance of the same genera (Fig. 1D) without any changes in alpha diversities across the developmental growth stages within each variety (Mondial: R = 0.004; p = 0.464 and Sifra: R= -0.235 and p = 0.836) (Fig. 2C). Similar to our observations, Pfeiffer et al., 2017 observed less diversity when the effects of plant development on bacterial diversity were investigated. This contrasts with reports on bacterial diversity changes particularly during the early plant stages and flower development [34, 37]. Community differences were observed only when varieties were compared as identified in our PCA (Fig. 2A). Phylogenetic clustering into 2 clades was observed (Fig. 2C) except for samples S3, S6 and S9 being phylogenetically closer to the Mondial bacterial communities. This supports a potato varietal influence on soil bacteria as previously reported in other studies [62]–[64]. Αn analysis of α-diversities showed that Mondial soils were richer than Sifra soils (Fig 3. A). Similar results were obtained by the Shannon index. An identification of taxa significantly contributing to these differences identified Mondial soils as rich in carbon degrading bacterial communities (Nocardioides iriomotensis strain IR27-S3 among others) while Sifra was enriched with nitrogen reducing bacteria (g_Gaiella and g_Conexibacter among others) (Table 1). Other studies have identified P. fluorescens F_140,_ Bacillus PTCC-1020, Strenotrophomonas maltophilia among many to associate with the potato plant [28, 60, 61]. The difference in these species suggests a unique association between potato variety and bacterial species.

The soil environment is considered one of the main determinants of the microbiome. Our results show a preferential proliferation of bacterial communities based on each field’s soil characteristic. Sifra soils were characterized by soil nutrients and negatively correlated with soil pH while Mondial samples were more correlated with high pH and base saturation. This suggests that Mondial soils lacked an abundance of nutrients whilst Sifra soils had low pH which ultimately reduces nutrient concentrations. Our study revealed pH as the environmental factor best determining bacterial communities between Sifra and Mondial (*p=0.001*). Sifra is described as a potato variety requiring low nitrogen requirement [67]. Despite this it is a plant that grows high with much foliage when it receives the right amount of N. Therefore, the heavy fertilization that occurs in potato production may have led to soil leaching and nitrogen deficiency. This would have ultimately led to decreased soil pH. *Gaiellea* from the Actinobacteria phylum have been implicated in nitrogen reduction [50] and the presence of numerous OTUs of these species in this study suggests a PGPR role that may have allowed N provision towards Sifra’s growth requirements (Table 1). Proliferation of these species would be directly dependent on Sifra’s exudates which signal PGPR assistance in N provision [39]. Unlike Sifra, Mondial is a high nitrogen input variety [59]. The ability of this variety to utilize more soil N may account for its higher pH compared to Sifra. The negative correlation of Mondial soils with nutrients suggests it as requiring more soil nutrients hence the presence of diverse PGPRs involved in nitrogen and carbon degradation.

An investigation of bacterial species physically attached to arbuscular mycorrhiza identified in the 2 potato fields revealed the presence of 4 bacterial genera, *Sphingomonas*, *Agrobacteria*, *Calidifontibacillus* and Bacillus. MHB in Mondial soils included *Sphingobium* and *Agrobacterium* MHB while Sifra AM fungi were associated with *Bacillus* and *Calidifontibacillus*. These genera have been historically documented to have MHB properties [61, 62]. Other bacteria such as Pseudomonas have been reported attached to AM fungi in potato soils [66]. An investigation of their functional aspects identified *Agrobacterium fabrum* strain C58 and *Sphingobium yanoikuyae* strain NBRC 15102 as capable of solubilization of complex phosphates, IAA, cellulase and chitinase production. Similarly, the MHB bacillus species had the same functionalities varying in their chitinase, protease and IAA producing capabilities (Fig. 5). The ability to produce chitinase, cellulase and protease also suggests their role in AM fungi hyphal growth, spore fitness and germination [68]. MHB have also been reported to have PGP properties. This concurs with the production of phosphatase and cellulase which suggests these species as assisting plants in nutrient uptake. Therefore, the beneficial effect of MHB are not always limited to AM fungal functionality but also potato growth and development [68].

In conclusion, the results from our study have identified core bacterial taxa that support Sifra and Mondial growth and development. These taxa are greatly influenced by potato genotype and prevailing soil environmental conditions. Therefore, in low soil pH, species belonging to the genera *Sphingomonas*, *f_Conexibacteraceae*, *f_Chthoniobacteraceae*, *Nocardioides*, *Solirubrobacter*, *Pseudolabrys* and *KD4_96_ge* are best suited in providing Sifra and Mondial plant growth promoting properties. Addition of MHB belonging to the *Sphingomonas* and *Bacillus* genera would be beneficial in supporting plant growth and AM fungi functionality. As bacterial communities vary with ecology and potato cultivars, an identification of core microbiomes is necessary across various ecological locations of the country to introduce sustainable potato production.

## 6.0 Acknowledgements

The authors would like to thank Dr Gwyneth Matcher from the South African Institute of Biodiversity at Rhodes University, for her technical assistance.

