## Supplementary data. Fig. 1 S1 for "Bacterial Communities and Soil Interactions in Sifra and Mondial Potatoes: Pathways to Sustainable Potato Production"

**8.0 Supporting information**

Supplementary material

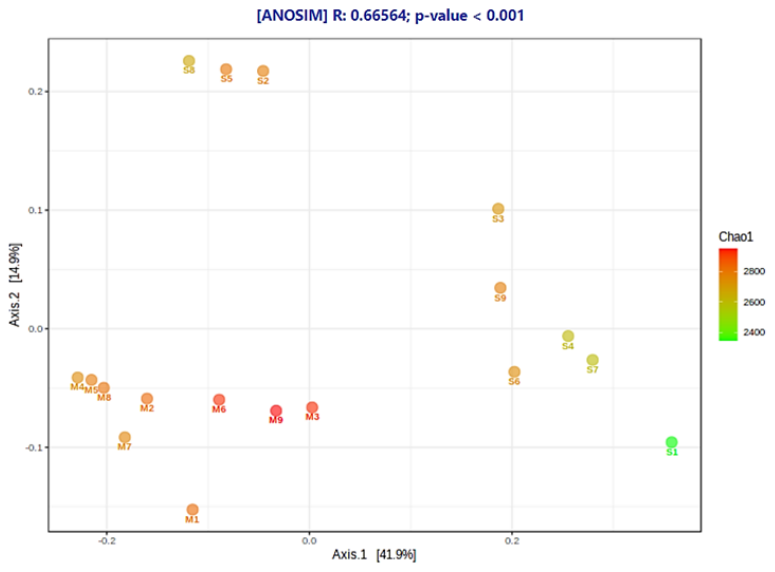

Fig. S1. Chao1 diversity differences when variety and growth stages are considered. Green denotes less diversity with red indicating high diversity. Mondial’s M3, M6 and M9 have the highest bacterial diversity.

Table S1

National Botanical Research Institute’s Phosphate growth medium (NBRIP)

| **Component** | **g/L** |
| --- | --- |
| Glucose | 100 |
| Ca_3_(PO_4_)_2_ | 5.0 |
| MgCl_2_.6H_2_O | 5.0 |
| MgSO_4_.7H_2_O | 0.25 |
| KCl | 0.2 |
| (NH4)_2_SO_4_ | 0.1 |
| Agar | 15.0 |
| Bromophenol Blue | 0.025 |

Autoclave at 121^0^C for 15 mins and cooled prior to dispensing into petri dishes.

Table S2.

**Chitin media**

| **Component** | **g/L** |
| --- | --- |
| Chitin from shrimp shell | 0.2 |
| Yeast extract | 5.0 |
| MgSO_4_ | 0.5 |
| Sodium nitrate | 2 |
| KCl | 0.5 |
| FeSO_4_ | Pinch |
| K_2_HPO_4_ | 1 |
| Agar | 20 |

pH to 6.0 and autoclave at 121^0^C for 15 mins and cooled prior to dispensing into petri dishes

Table S3.

**Carboxymethylcellulose media**

| **Component** | **g/L** |
| --- | --- |
| NaNO_3_ | 2.0 |
| K_2_HPO_4_ | 1.0 |
| MgSO_4_ | 0.5 |
| KCl | 0.5 |
| CMC Na Salt | 2.0 |
| Peptone | 0.2 |
| Agar | 17.0 |

Autoclave at 1210C for 15 mins and cooled prior to dispensing into petri dishes.
